# Prey movement shapes the acquisition of predator expertise in a virtual bi-trophic system

**DOI:** 10.1101/2024.11.15.621573

**Authors:** Maxime Fraser Franco, Francesca Santostefano, Julien G. A. Martin, Clint D. Kelly, Pierre-Olivier Montiglio

## Abstract

The acquisition of expertise is crucial for predators to be successful hunters. To achieve this, predators must hone their skills and gain knowledge through repeated and extensive practice. On the other hand, prey may hinder the acquisition of predator expertise by employing antipredator tactics to evade detection and pursuit. However, empirical evidence on how predators acquire expertise through repeated encounters with their prey remains limited, largely due to the challenges of monitoring direct interactions in the wild. Here, we use a virtual predator-prey system (the game *Dead by Daylight*) to investigate how experience shapes individual and population hunting success in human predators across repeated interactions with their prey. We show that predators optimized prey consumption as they gained experience, indicating that they acquired expertise through extensive practice. At the population-level, we found that faster prey impaired the acquisition of expertise by reducing hunting success. Prey speed was also an important mediator of this relationship at the individual level, driving differences among predators in the acquisition of expertise. Our study outlines how prey antipredator behaviour can mediate the acquisition of expertise in predator populations.

## INTRODUCTION

Predation is a fundamental biological process that acts as an agent of evolutionary change and shapes community structure (Abrams 2000; Wirsing et al. 2021). By consuming prey, predators regulate prey populations, limit the spread of diseases, and mediate energy flow across trophic levels (Hairston Jr. and Hairston Sr. 1993; Ripple et al. 2014; Schmitz 2017). These processes emerge from interactions between individual predators and their prey, lying at the core of key concepts such as predation risk and prey selection(Pettorelli et al. 2015; LaBarge et al. 2024).

Predators also respond to interactions with their prey, increasing their effectiveness by adapting their strategy and choosing suitable habitats to hunt (Quevedo, Svanbäck, and Eklöv 2009; Schmitz 2017). Central theory suggests that these responses by predators may result from acquiring expertise through practicing and learning the proper skills to locate, select, and capture their prey (Woo et al. 2008; Wooster et al. 2023; LaBarge et al. 2024). Hence, the role of expertise remains a missing but essential link to assess of how individual characteristics shape prey consumption during predator-prey interactions.

Expertise can be defined as the characteristics, skills, and knowledge that provide individuals with the ability to outperform novices on complex tasks (Dukas 2017). Empirical studies on human and non-human hunters show that individuals optimize foraging efficiency (e.g. search and handling times, return rates) by associative learning, by developing search images, or by exploiting cues from their prey and their environment (Edwards and Jackson 1994; Morse 2000; MacDonald 2007; Reid, Seebacher, and Ward 2010; Wilson-Rankin 2015). Through these processes, expert predators should have greater knowledge, better energy management, and acute motor skills to increase their chances of locating and capturing prey (Dukas 2019). Therefore, differences among predators in their capacity to acquire expertise throughout their lifetime may underlie differences in hunting success. This is an important topic of study since predator cognition can maintain behavioural variation, shape prey phenotypic composition, and destabilize predator-prey systems (Kondoh 2010; Skelhorn and Rowe 2016; Kikuchi and Simon 2023).

Prey use antipredator tactics such as camouflage to avoid detection and rapid escapes to evade capture (Walker et al. 2005; Kelley and Magurran 2011; Herbert-Read et al. 2017). These strategies are hypothesized to drive differences in prey consumption among predators by disrupting the acquisition of hunting expertise (Wooster et al. 2023). Experimental studies have shown that certain camouflage tactics of prey can impair expertise acquisition in humans and birds (Stevens et al. 2012; Troscianko et al. 2013). For example, Troscianko, Skelhorn, and Stevens (2018) found that disruptive colouration interfered with human subjects’ ability to form search images, hindering improvements in detection times over repeated attempts. Thus, antipredator behaviour may also hold the potential to hinder the acquisition of predator expertise. For instance, some predators may be limited in their capacity to develop the necessary attributes (e.g. physical, physiological, neurological) for fast-paced hunting, which could impair their acquisition of hunting expertise. To our knowledge, there is no empirical evidence showing links between prey antipredator behaviour and the acquisition of expertise in human and nonhuman predators. This represents a significant gap in our understanding of predator-prey interactions, as individual variation in predator expertise could lead to shifts in foraging behaviour and define whether predators select prey based on their frequency or their traits (Allen et al. 1988; Ishii and Shimada 2012; Wooster et al. 2023).

A recurring challenge impeding research on predator-prey behavioural interactions is the need to collect data simultaneously on both the predator and the prey. Here we mitigate these challenges by using *Dead by Daylight* as our study system, a videogame where four prey players must forage for resources while avoiding predation by a fifth player. Similar to agent-based simulations, *Dead by Daylight* provides controlled virtual environments to test ecological hypotheses (see Lymbery, Webber, and Didham 2023 for an example with *Age of Empires II*), but with the advantage of having real players that interact in the virtual space. In this game, the predator population comprises individuals that either ambush or hunt at high speeds (i.e. mean movement speed along a slow-fast continuum), and their success is driven by the movement of the prey (Fraser Franco et al. 2022). The prey can increase their chances of survival by cooperating and moving fast to escape the predator (Céré, Montiglio, and Kelly 2021; Fraser Franco et al. 2022; Santostefano, Fraser Franco, and Montiglio 2024). The game also elicits natural reactions in players such as freezing when predation is imminent (M.F.F., personal observations), corroborating another virtual ecological study showing that predation drives individual variation in risk perception (Beauchamp 2020). These observations outline how ecological phenomena can emerge from human interactions in virtual systems with fixed rules (Brosnan and Postma 2017; Kasumovic, Blake, and Denson 2017). Videogames also generate large volumes of data on thousands of interacting players throughout their lifetime in the game under realistic, controlled, and repeatable ecological scenarios. Hence, *Dead by Daylight* allows us to tackle fundamental questions about the role of antipredator behaviour and experience in predator-prey interactions.

In this study, we assess how repeated encounters with prey shapes predator hunting success using data from players in *Dead by Daylight*. We quantify expertise acquisition as the relationship between hunting success (i.e. the probability of capturing all prey) and repeated experience (i.e. cumulated atempts). First, we test the hypothesis that predator success will increase with experience up to some point at which it will stabilize (Dukas 2019). However, we expect this pattern to change depending on the movement of the prey encountered. We hypothesize that prey will influence expertise acquisition, and predict that faster prey will reduce the gain in expertise. Therefore, we also investigate how prey movement influences expertise acquisition at the individual level. If prey speed does not influence hunting success, we predict that the gain in expertise will be similar among individuals. Alternatively, if prey speed influences hunting success, then the acquisition of expertise will vary among individuals.

## MATERIALS AND METHODS

### Study system

*Dead by Daylight* is a survival asymmetric (i.e. gameplay mechanics differ between two groups) multiplayer online game developed by Behaviour Interactive Inc., in which players can play either as predators or prey. Every match includes only one predator and four prey. The objective of the predator is to hunt and capture the prey, and the objective of the prey is to search for resources while avoiding the predator. The resources are in the form of power generators that, once all activated, will enable the prey to escape through one of two exit doors. A skill-based matchmaking algorithm determines the composition of the predator and prey group in a match. A match ends when the predator kills all the prey available (i.e. that have not escaped), or when the last remaining prey escapes the virtual environment.

Before the start of a match, players (predator or prey) can choose an avatar with unique abilities that encourage specific play styles (e.g. bold vs cautious prey, or ambush vs roaming predator). During our study period, the game offered 23 predator avatars. The virtual environments comprise fixed and procedurally generated habitat components, such as vegetation, mazes, and buildings. Some of these environments are larger than others, with varying structural complexity. However, predators display only minimal changes in behaviour and hunting success across these environments (Fraser Franco et al. 2022). There were 35 virtual game environments available for play during the study period. Details on the basic characteristics of predator avatars are available at https://deadbydaylight.fandom.com/wiki/Killers. Details on the size and structure of the different virtual environments are available at https://dbdmaps.com/ and https://deadbydaylight.fandom.com/wiki/Realms.

### Data collection

Behaviour Interactive Inc. provided data that spanned six months of gameplay recorded for every player from 2020-12-01 to 2021-06-01. We analyzed only matches where players did not know each other. We filtered any matches where players were inactive, such as when mean distances travelled per second (i.e. speed) were equal to, or very close to, zero. Moreover, we used our knowledge of the game to remove any matches where players were potentially hacking, or not playing the game as intended.

We sampled players that played 300 matches or more and monitored all their matches from the first to a maximum of 500 matches. We recognize that we could have introduced a bias by retaining only those individuals, as they might already be seasoned video game enthusiasts and exhibit expert-level performance in their early matches in *Dead by Daylight*. Thus, we verified that our sample was not biased by comparing a random sample of players that played either 20 to 50 matches, 51 to 100 matches, or 101 to 300 matches during the same timeframe as our sampled population. We then took the first 20 matches played by these players, including those from our sampled population, and compared their median hunting success using a Bayesian hierarchical linear model. We found that all four groups had similar success as predators (Appendix 1: Table S1 and Figure S1), suggesting an absence of bias due to data sampling.

Our population consists of 253 players who played as the predator, with a total record of 100 412 matches. The predator-players’ experience varied between 301 and 500 matches played. These matches lasted between 3 and 70 min (mean = 11 min). The following information is collected and reported for every match : the player’s anonymous ID, its avatar (i.e. the predator character chosen with its specific skill-gameplay mechanics), the game environment, the predator-player’s experience, the mean speed of the groups of prey that the predator player encountered, and the mean rank of the prey encountered (a proxy for prey skill). The ranking system in *Dead by Daylight* was implemented by the company to pair players in a match based on their skill (https://deadbydaylight.fandom.com/wiki/Rank), and failing to account for it would prevent us from detecting a change in the predator’s foraging success with experience.

We analyzed the mean speed of the prey group encountered by the predator. We measured the prey’s speed as the mean travel speed of the four individual prey in a match (mean = 2.40 ± 0.32 m/s). We defined hunting success as the number of prey consumed during the match (min = 0, max = 4). Lastly, we defined the predator’s cumulative experience as the number of matches played as the predator prior to the match being monitored. For example, the first match of a player would have a cumulative experience value of 0, while the tenth match would have a value of 9. We did not account for matches where predators played as the prey.

### Data analyses

#### Model specification

We tested how predators developed their expertise by computing five Bayesian generalized additive mixed models (GAMM) with thin-plate regression splines, all of which estimated the relationship between hunting success (i.e. number of prey consumed) and the predators’ cumulative experience (i.e. number of matches played before the current match). We parametrized the models following the method of Pedersen et al. (2019). The first model (I) was the simplest, with a common global smoothing function and random intercepts for the predator ID. In this model, we assume that predators have the same acquisition of expertise, with the model estimating a trend for the average individual (i.e. global smoother). The second model (II) included varying individual smoothers for the predator ID. Here, we assume that individual predators share a similar relationship between success and experience, but that this relationship can vary among them (e.g. predator 1 has a steeper curve than predator 2). This enabled us to test whether predators differed in the development of their expertise. In the third model (III), we kept the individual smoothers for the predators, but removed the global smoother. This model assumes that predators do not share a common relationship between success and experience. The fourth (IV) and fifth (V) models were expansions of the third and second models respectively, where we included the prey speed to assess its effect on the relationship between success and experience. We included the standardized match duration and prey rank as covariates in all five models.

We computed the five models using a modified version of the beta-binomial distribution. Hunting success was estimated as the probability of consuming the four prey (*μ*_*i*_), drawn from a Beta distribution (*Beta*(*μ*_*i*_, *ϕ*)) with mean (*μ* ∈ [0,1]) and precision (*ϕ* > 0) parameters. We used a logit link function to estimate *μ*_*i*_ where 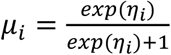 and *η*_*i*_ is the linear predictor, while the precision parameter (*ϕ*) was estimated with an identity link. We used ten basis functions (K = 10) for the models to estimate the relationship between hunting success and experience. We assumed that the random intercepts for the predator ID (*id*) followed a Gaussian distribution with estimated standard deviation 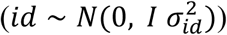). We fitted all models in R (version 4.1.2) using Hamiltonian Monte Carlo (HMC) sampling with the package “brms” version 2.16.3 (Bürkner 2017), an R front-end for the STAN software (Stan Development Team 2023), and “cmdstanr” version 0.4.0 (Gabry and Češnovar 2021) as the back-end for parameter estimation (cmdstan installation version 2.28.2). For further details, please consult the GitHub repository of this project (https://github.com/quantitative-ecologist/predator-expertise).

We used weakly informative Gaussian priors for the intercept (*N*(0, 0.5^2^)) and the global trend of cumulative experience (*N*(0, 2^2^))). Following Fraser Franco et al. (2022), we defined a positive Gaussian prior on the precision parameter (*N*(2, 0.5^2^)), a positive Gaussian prior (*N*(1, 0.5^2^))) on the game duration because longer trials lead to greater success, and a negative Gaussian prior on prey speed (*N*(−1, 0.5^2^)) because encountering faster prey is associated with lower success in this system. We employed weakly informative half-Gaussian priors on all the standard deviation parameters (*N*(0, 0.5^2^)). We compared the predictive accuracy of all five models using approximate leave-one-out cross-validation with Pareto-smoothed importance sampling (Vehtari, Gelman, and Gabry 2017; Piironen and Vehtari 2017; Vehtari et al. 2022).

#### Parameter sampling settings

We parametrized the GAMMs to run four chains. We ran 2500 iterations with a thinning set to eight for model I (see Table 1), and 1500 iterations with a thinning set to four for the remaining additive models (models II to V). We set the first 500 iterations of each model as warm-ups. For each model, we obtained 1000 posterior samples per parameter. We assessed the convergence of the chains using trace plots, R-hat diagnostics with a threshold of <1.01, and effective sample sizes (ESS) with a threshold of >100 (Vehtari et al. 2021). We also performed posterior predictive checks which showed an adequate fit of the models.

**Table 1.** Leave-one-out cross-validation table of the five GAMMs relating hunting success to predator experience.

| model | elpd difference | sd difference | elpd loo value | elpd loo standard error |
| --- | --- | --- | --- | --- |
| (V) predator xp + ID smoothers + prey rank + prey speed | 0.00 | 0.00 | -136 123.69 | 201.04 |
| (IV) ID smoothers + prey rank + prey speed | -562.90 | 23.59 | -136 686.59 | 202.06 |
| (III) ID smoothers + prey rank | -5 717.54 | 107.99 | -141 841.22 | 184.27 |
| (II) predator xp + ID smoothers + prey rank | -8 536.39 | 129.62 | -144 660.08 | 197.49 |
| (I) predator xp + ID intercepts + prey rank | -8 593.08 | 131.73 | -144 716.77 | 187.16 |

<sup>a</sup> 'elpd' refers to the expected log pointwise density and is the value chosen to select the best model.
<sup>b</sup> 'xp' is an acronym for experience

#### Hypothesis testing

We tested the hypothesis that antipredator behaviour impairs the acquisition of expertise at the population level by visually comparing the global trends of model II (not controlling for prey speed) and model V (controlling for prey speed) relating hunting success to cumulated experience (Figure 1). At the individual level, we tested our hypothesis that antipredator behaviour generates differences among predators in expertise acquisition by comparing individual-level parameters of model II and model V. Specifically, we compared the standard deviations of 1) the random intercepts (i.e. mean differences in hunting success), 2) the random slopes (i.e. linear component relating hunting success with experience), and 3) the curve wiggliness (i.e. nonlinear component relating hunting success with experience).

**Figure 1.**
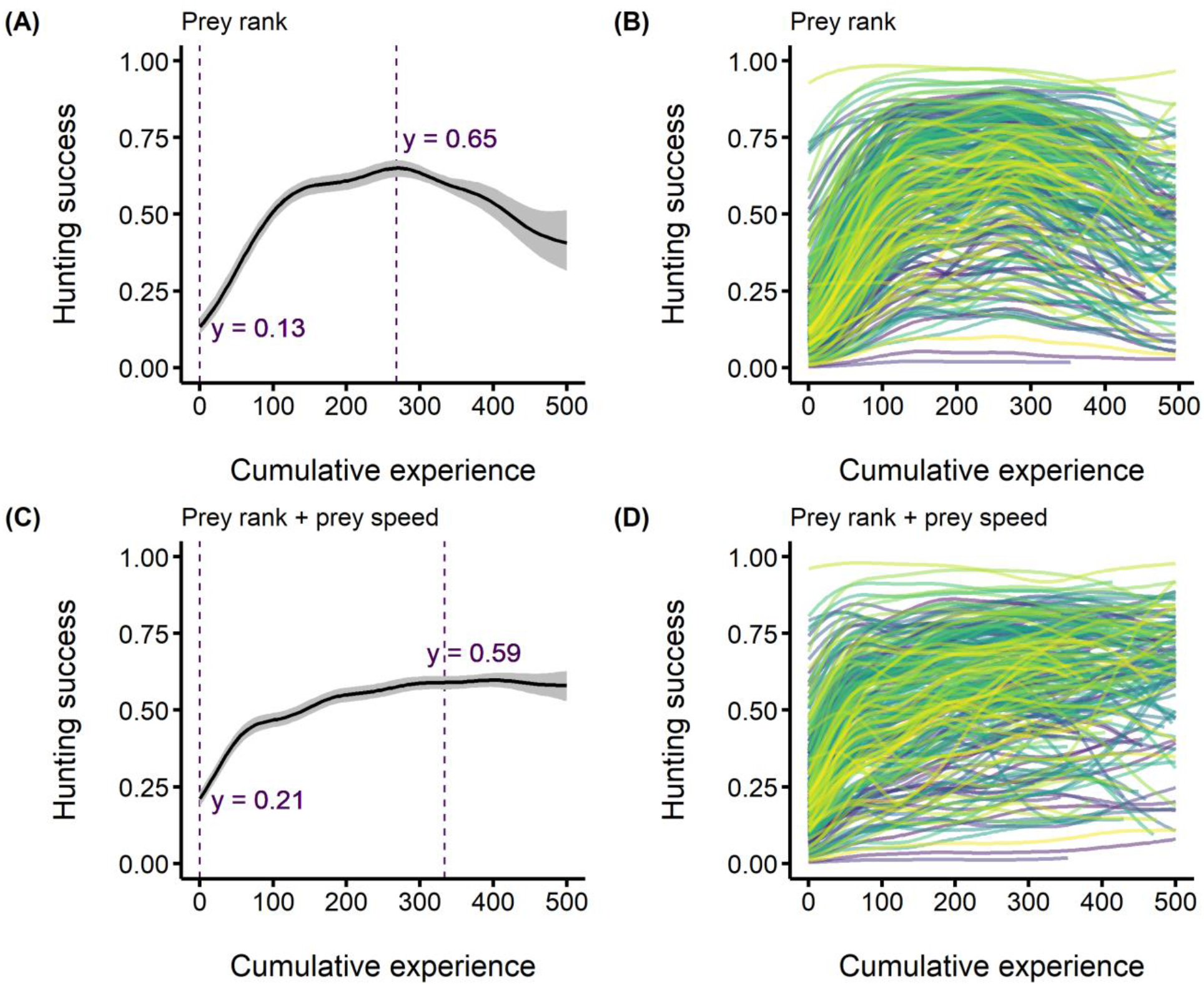
Median posterior predictions of the acquisition of predator hunting expertise. The predators’ hunting success (i.e. the probability of consuming the four prey) is on the y-axis, and the predators’ cumulative experience (i.e. the number of matches played before each observation) is on the x-axis. Panels A and C show the acquisition of expertise for the average individual with the vertical dashed lines on the left representing the lowest predicted values. For panel A, the right-side vertical dashed line shows the highest predicted success. For panel C, the right-side dashed line represents the point on the curve where success was optimized, which we calculated using the finite differences method to obtain the first derivative of the predicted values. Panels B and D show among individual differences in the acquisition of expertise, with each curve representing an individual predator. (A-B) model II where we control for the prey rank (C-D) model V where we control for the prey rank and the speed of the prey group.

## RESULTS

### Acquisition of expertise at the population level

Out of all five GAMM models, the two that accounted for the prey group’s rank and speed were the best at predicting the data with similar predictive accuracies (Table 1). Models in which prey rank was not accounted for resulted in no change in hunting success with experience for the average individual (i.e. no gain in expertise, results not shown). Accounting for the prey rank resulted in a concave-shaped relationship (model II), with the highest success ranging between ∼200 and ∼300 matches (Figure 1A).

We found strong evidence of a negative relationship between hunting success and prey speed (Figure S2). As predicted, the effect of experience on hunting success for the average individual followed a diminishing returns curve when controlling for prey speed (model V), with predators optimizing their success after playing ∼300 matches (Figure 1C). The curve shows a 38% increase in the probability of consuming all prey for the average individual between the first and the ∼330^*th*^ match, where success reached a plateau (Figure 1C).

### Acquisition of expertise at the individual level

Prey speed did not influence among-individual differences in average hunting success as the posterior distributions of the standard deviations of individual intercepts were almost completely overlapping (Figure 2, median = 2.21 vs median = 2.19). However, individuals differed in the acquisition of their hunting expertise (Figure 1B-D). We found strong evidence that the speed of the prey mediated among-individual differences in the linear relationship between success and experience, as there were substantial differences in the standard deviations of the individual slopes between the two models (Figure 2, median = 9.72 vs median = 3.35). Differences among individuals in the direction of the linear relationship between success and experience were 2.9 times lower when we removed the effect of prey speed (i.e. accounting for it in the model).

**Figure 2.**
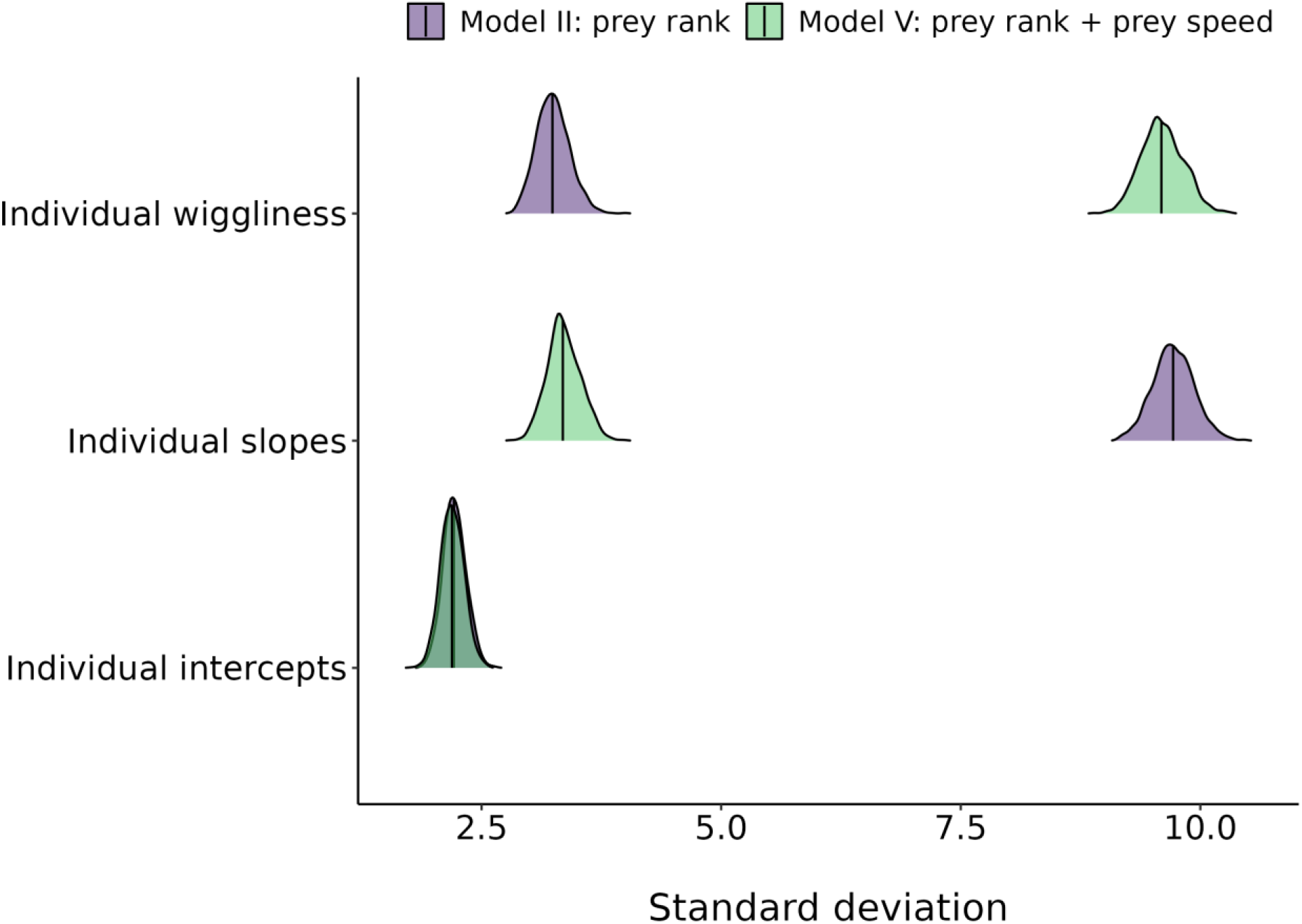
Posterior distributions of the standard deviation of individual-level parameters estimated by the GAMM. The parameters are displayed on the y axis, and their standard deviation are displayed on the x axis. The intercept and slope standard deviations refer to the linear components of the estimated relationship between hunting success and cumulative experience. The standard deviation of the wiggliness parameter refers to the shape of the curves (i.e. nonlinear component). The vertical lines are the medians of the posterior distributions. For each panel, the light-coloured distributions (i.e. green) are for the model with a shared trend where we did not account for prey speed (model II), and the darker-coloured distributions(i.e. purple) are for the model with a shared trend where we accounted for prey speed (model IV).

We also found strong evidence that the speed of the prey mediated the form of the relationship between hunting success and experience at the individual level. We detected large differences in the standard deviations of the wiggliness (Figure 2, median = 3.24 vs median = 9.59). The lower standard deviation for the model where we accounted for prey speed (model V) suggests that the form of the relationship between success and experience was more similar among individuals.

## DISCUSSION

Using a virtual predator-prey system where we monitored predator hunting success across experience, we provide rare empirical support for the hypothesis that prey antipredator behaviour can impair the acquisition of hunting expertise. We show that the predator population increased its hunting success with experience. We found, however, important differences among individuals in expertise acquisition that were related to the speed of the prey encountered.

Our results suggest that predators hone their hunting expertise through extensive practice. The predator population displayed an asymptotic relationship between experience and success, wherein initial gains in success were significant but gradually stabilized as experience accumulated. These observations are consistent with empirical studies of expertise in both humans and nonhuman animals (reviewed in Dukas 2019). However, prey speed played a crucial role in shaping this pattern at the population level as encounters with faster prey resulted in lower hunting success (Figure S2). We previously showed in *Dead by Daylight* that faster movement is effective for prey to evade predation (Fraser Franco et al. 2022), in agreement with studies in other animals (Walker et al. 2005; Kelley and Magurran 2011; Martin et al. 2022). We suspect that experienced prey may increasingly rely on this strategy, which could explain why the relationship between hunting success and experience was concave when we did not control for prey speed (model II, Figure 1A). Thus, our results suggest that predators can gain expertise and maintain success when they encounter prey that move at speeds lower than or closer to the population-average.

Prey speed also mediated differences among predator players in the acquisition of expertise, suggesting that individual predators varied in their capacity to adjust to challenging prey. Animals are expected to have limited attention spans, which restricts diet choice and the formation of search images (Dukas and Kamil 2001). Hunting faster prey demands specialized cognitive abilities and coordination that are energetically costly (Kelley and Magurran 2011). Thus, predators that failed to develop counter-strategies for detecting or chasing faster prey were likely at a disadvantage. Parallel observations have been outlined in studies of prey camouflage strategies. For example, Troscianko, Skelhorn, and Stevens (2018) showed in a computer experiment involving humans that disruptive camouflage was efficient at preventing the acquisition of expertise during search image formation. Human subjects exposed to a restricted set of strategies were also less efficient compared to those exposed to a variety of strategies. Therefore, our observations show that prey antipredator behaviour can also impair predator expertise acquisition.

Despite adjusting for the prey’s speed (i.e. model V), noticeable differences in expertise acquisition among predators persited. One possible explanation is that the predators’ hunting tactics may indirectly shape their own expertise through changes in prey behaviour. Predators tend to specialize as cursorial or ambush hunters in *Dead by Daylight* (Fraser Franco et al. 2022). Consequently, those employing a cursorial tactic may push prey to move faster, which, in turn, may hinder their own expertise acquisition due to the increased difficulty of hunting such prey.

An alternative explanation is that longer time intervals between hunting events are hypothesized to hinder or delay the acquisition of expertise because individuals may forget information when delays are longer (Endler 1991; Wright et al. 2022). For example, a predator that played 300 matches in six months might forget critical information related to prey detection or escape patterns compared to one that played 300 matches in six days. While there is unequivoqual evidence that many predator species can learn quickly how to be efficient hunters, the role of the frequency of interactions remains unclear (Wooster et al. 2023). Therefore, investigating the impact of such time lags in future analyses may reveal important insights into the outcome of predator-prey interactions. Another potential reason for the persistent differences among individuals is that neither the predator nor prey players’ lives are at stake in the game. As a result, emerging patterns may be driven more by the players’ motivation to win rather than “true” survival. For example, some players could experiment with the game out of boredom, which could also contribute in shaping how expertise is honed in this particular system.

## Conclusions

We found support of our hypothesis that prey antipredator behaviour drives individual differences in expertise acquisition in a human predator population in the game *Dead by Daylight*. Future analyses should investigate how antipredator tactics developp with experience, as it may reveal important insights on the eco-evolutionary dynamics of predator-prey interactions. Our study demonstrating that prey antipredator behaviour can impair the acquisition of hunting expertise adds to a growing body of research showing how virtual systems can be used to test hypotheses on ecological interactions (Beauchamp 2020; Céré, Montiglio, and Kelly 2021; Fraser Franco et al. 2022; Lymbery, Webber, and Didham 2023; Santostefano, Fraser Franco, and Montiglio 2024). We therefore hope that our study will inspire more collaborations between scientists and the videogame industry to tackle fundamental questions in ecology.

