## Appendix for "Prey movement shapes the acquisition of predator expertise in a virtual bi-trophic system"

Table S1. Fixed effects table comparing the success of random groups of players with different amounts of matches played to the group presented in the main text.

| Parameter | Estimate | lower 89% CI | upper 89% CI |
|---|---|---|---|
| game duration | 0.56 | 0.55 | 0.58 |
| cumulative experience | 0.11 | 0.09 | 0.12 |
| group 1 | 0.01 | -0.06 | 0.08 |
| group 2 | 0.05 | -0.03 | 0.14 |
| group 3 | 0.09 | 0.01 | 0.16 |
| group 4 | -0.03 | -0.15 | 0.09 |

[a] Group 1: <50 matches, Group 2: between 50 and 99 matches, Group 3: between 100 and 299 matches, Group 4: > 299 (i.e. group in the main text)

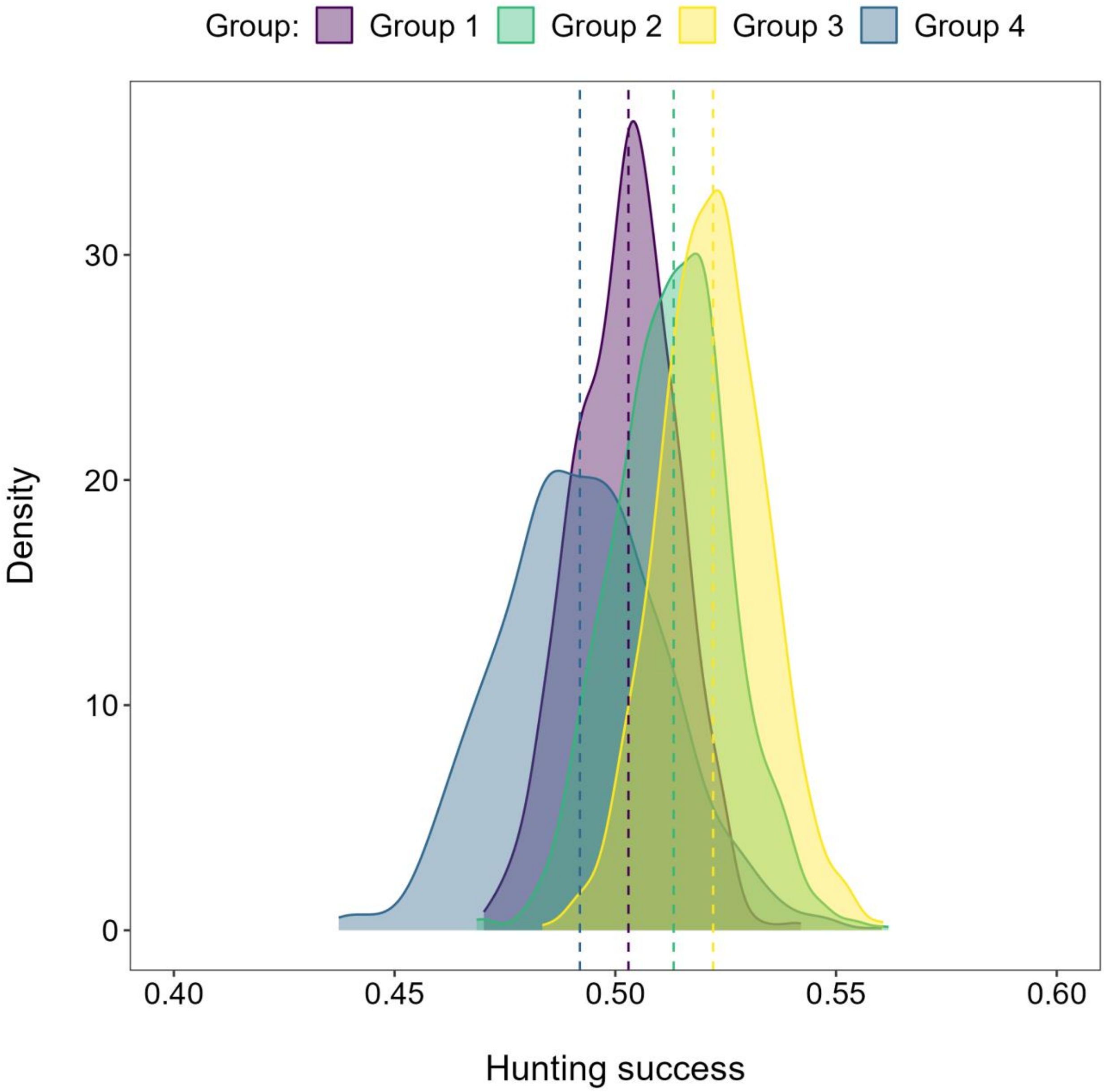


Figure S1. Posterior distributions of the median hunting success per group of players. The values were back transformed to probability scale and can be interpreted as the probability of capturing the four prey. Group 1: <50 matches, Group 2: between 50 and 99 matches, Group 3: between 100 and 299 matches, Group 4: > 299 (i.e. group in the main text).

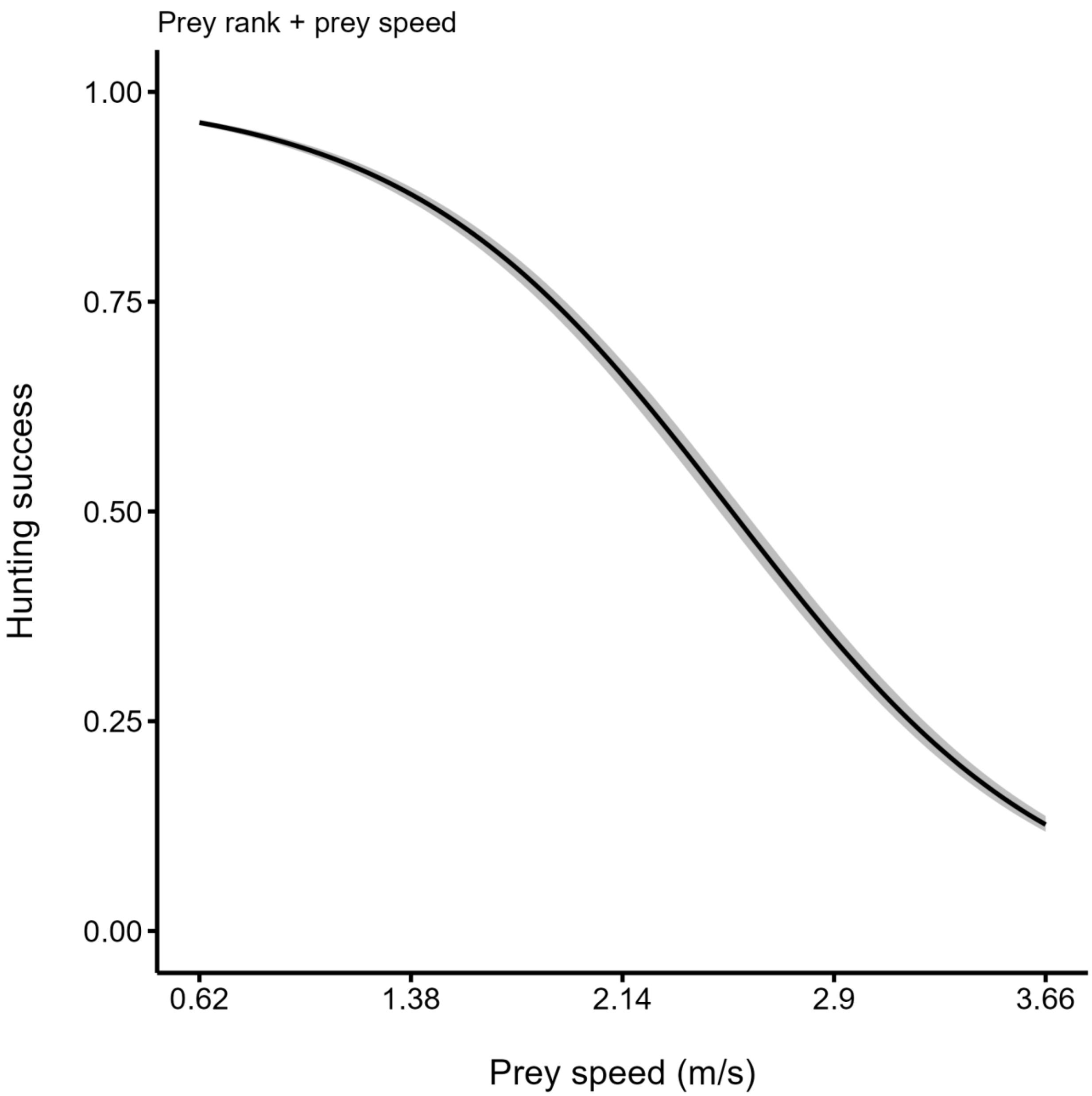


Figure S2. Median posterior predictions of the relationship between hunting success (i.e. the probability of capturing the four prey) and the average speed of the prey group. The black line represents the median posterior predictions, and the gray band the the 89% compatibility interval. The predictions are from the model with the highest predictive accuracy (see Table 1 in main text).
